# Habitat structure drives the evolution of aerial displays in birds

**DOI:** 10.1101/480053

**Authors:** João C. T. Menezes, Eduardo S. A. Santos

## Abstract

Physical properties of the environment may shape signalling traits by determining how effective the signals are in affecting the behaviour of other individuals. Evidence abounds of signalling environment driving the evolution of colours and sounds, but little is known about its influence upon gestural displays. Here, we performed a continent-wide phylogenetic comparative analysis to test the hypothesis that habitat structure drives the evolution of aerial sexual displays in passerine birds. We found that aerial displays are seven times more likely to evolve in open habitats than in forests, likely as a result of physical properties that allow aerial displays to transmit more broadly in open habitats. Our results provide an emblematic example of how environmental factors may help predict the direction of evolution of otherwise unpredictable sexual traits. The broader range of aerial displays in open habitats may also mean that females can sample more males, potentially leading to more intense sexual selection over open-habitat, aerial-displaying males.

---

Communication in animals occurs through the emission and reception of signals — acts or structures that, by definition, have been selected because they affect the behaviour of other organisms^1,2^. How effectively a particular trait affects the behaviour of another organism, however, depends upon the physical properties of the environment through which the signal is transmitted^3^. For instance, females of three-spined sticklebacks (*Gastosteus aculeatus*) prefer mates that have a brighter-red coloration^4^. Red coloration in males, thus, is a signal: it has been selected because it affects how females choose a mate. However, under green artificial light, red coloration is not transmitted as effectively and thus females show no preference^4^. If a hypothetical population of sticklebacks had lived under green lighting conditions all along their evolutionary time, red coloration would most likely not be selected as a signal of male quality. Such influence of environmental properties upon both ends of communication systems – signalling trait and sensory tuning to receive it – is the key feature underlying the sensory drive hypothesis^3,5,6^. As a consequence, we can expect the same characteristic to be positively selected in some environments, but not in others. Accordingly, the structure of many signals has been found to be habitat-dependent in animals that use colours, sounds, ornaments or vibration to communicate (reviewed in ^3,7^). However, empirical evidence of sensory drive upon gestural (motion-based) signals (henceforth, displays) is mostly restricted to individuals adjusting their behaviour to maximize the conspicuousness of the display – for example, by choosing an appropriate signalling site or timing^8–13^.

Various animals, from jellyfishes^14^and arthropods^15–17^to aquatic and terrestrial vertebrates^18–24^, use gestural displays as sexual signals. These displays are selected through intersexual mate choice or intrasexual competition mechanisms^25,26^and may, as any signal, be subject to selective pressures imposed by the signalling environment. Only recently, however, have researchers begun to look into the potential influence of the environment upon the structure of displays – rather than the choice of timing or site –, with two studies finding evidence of it^27,28^. Thus, we currently have tentative knowledge about the influence of the signalling environment on the evolution of gestural displays.

Sexual displays performed by passerine birds come in all forms: from the simple, static ‘bill-up’ posture of silver-beaked tanagers (Isler & Isler 1987 *in* ^29^) to the most complex, skilful dances performed by manakins^30^and birds-of-paradise^28^. Passerines also exhibit high variability in habitat preferences, having colonized environments as diverse as Arctic tundras, tropical rainforests and arid deserts^31^. These features make Passeriformes a promising group to investigate the potential role of environment in shaping displays. As we will argue, a particular component present on many of the sexual displays exhibited by passerines – the flight component of aerial displays – seems particularly likely to evolve under sensory drive. For example, flying above an open habitat (*vis-à-vis* underneath the forest canopy) during the display puts the signaller under more intense light^32^, and, as a consequence, the signal reaches a wider range of potential receivers, all else held constant^33^. Moreover, flying above an open habitat may result in less or no vegetation obstructing the signaller from the perspective of potential receivers. Flying inside the forest does not reduce visual obstacles between signaller and receivers and, in addition, the display is less likely to be perceived the farther signaller and receiver are because vegetation accumulates horizontally. Lastly, complex visual backgrounds make signals less consistently perceived^9^. The sky is as uniform as a background may be, while forest interior is a highly heterogeneous one.

Thus, although females may assess motor performance of males from a short distance in both habitats^26^, it is unlikely that aerial displays offer any advantage in a broader spatial scale in forests. In open habitats, a flight component may allow the display reach a broad range of potential receivers with more intensity, less visual obstruction, and more consistency. Here, we evaluated whether sensory drive influences the evolution of sexual displays by testing the hypothesis that aerial displays are more likely to evolve in passerine birds that inhabit open habitats than in those that inhabit forests. We did that by performing a continent-wide phylogenetic analysis of 469 species, which constitutes the most comprehensive test of sensory drive – phylogenetically and geographically – as far as we are aware. If our findings support this hypothesis, we would provide evidence to corroborate, for the first time, the pattern predicted for decades by researchers^34–36^, but never actually tested, that aerial-displaying species are more prevalent in open habitats than in forests.

## Results

We were able to collect data about sexual display and primary habitat structure for 469 (19.2%) species of New World passerines belonging to 41 families (Supplementary Data); the remaining species – for which data was not available – were excluded from analyses. Each species was classified as to the *presence* (1) or *absence* (0) of aerial sexual display, and as living primarily in *open habitats* (1) or *forests* (0). Among open-habitat passerines, 127 of 188 species (68%) were aerial displayers, whereas 48% (*N* =134) of the 281 forest species exhibited aerial display (Figure 1). Although this pattern is consistent with the common perception that aerial-displaying passerines are more prevalent in open habitats, it allows no inference as to whether habitat influences the evolution of aerial display.

**Figure 1.**
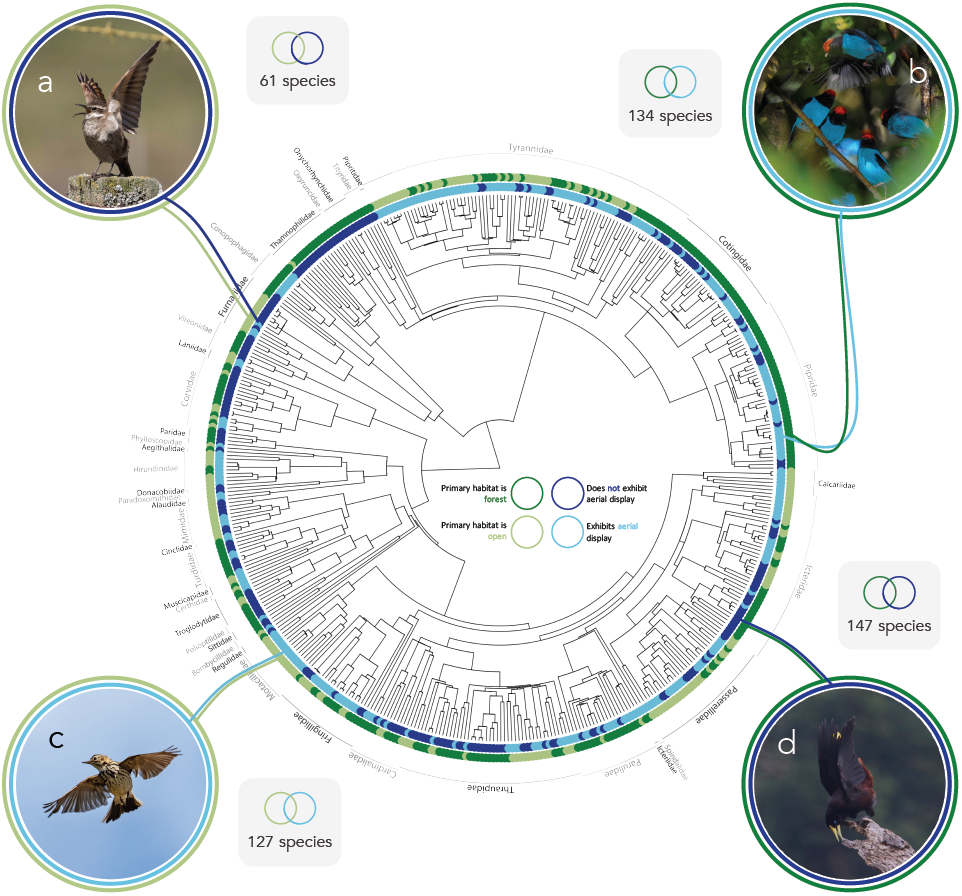
Phylogenetic distribution of display and habitat structure data (inner and outer circle, respectively), number of species (grey boxes) and examples of species in each category: (a) stout-billed cinclodes (*Cinclodes excelsior*) exhibit non-aerial display in open habitats (photo courtesy of Ken Chamberlain), (b) blue manakins (*Chiroxiphia caudata*) exhibit aerial display in forests (photo courtesy of João Quental), (c) meadow pipits (*Anthus pratensis*) exhibit aerial display in open habitats (photo courtesy of Kevin Hay), and (d) crested oropendolas (*Psarocolius decumanus*) exhibit non-aerial display in forests (photo by Gregory Smith, CC BY-SA 2.0). For the ancestral state estimation, see Figure S1.

To test the hypothesis that aerial displays are more likely to evolve in open-habitat passerines than forest ones, we conducted two different phylogenetic comparative analyses using species-level Passeriformes trees^37^. In the first analysis, we fitted phylogenetic logistic regression models (PLogReg^38^) to determine whether habitat structure influences the probability that a species exhibits aerial display, while accounting for phylogenetic dependence among species. We compared a null model (with no predictor variable) with one in which habitat structure was fit as a categorical predictor, and accounted for phylogenetic uncertainty by running each model with a sample of 1,000 topologies, and performed model choice using the Akaike information criterion (AIC). The model containing habitat structure as a predictor was selected in all of the 1,000 iterations, with a mean ΔAIC of 17.56 (min = 6.11, max = 46.23). Mean slope estimate for open habitat was 0.489 (95% CI: 0.485 to 0.492), with a mean *p-*value of 0.025 (≤ 0.05 in 92.8% of iterations). This indicates that the probability that a passerine species exhibits an aerial sexual display is greater in open habitats than in forests. Performing this analysis with a Bayesian (MCMC) approach yielded qualitatively similar results (see Supplementary Results). We performed an additional PLogReg to test whether forest stratum influences the probability that forest passerines exhibit aerial display, but found no clear effect (see Supplementary Results). This result does not support the idea that canopy stratum acts in a similar way as open habitats in terms of signal transmission properties.

Secondly, we fitted a series of Markov models^39,40^to test for correlated evolution between aerial display and habitat structure using the same set of species and trees. We designed four biologically plausible models: a) independent (habitat transitions not depending on aerial display state and vice-versa), b) correlated (each variable depending on the other), c) habitat-dependent (aerial display transitions depending on habitat state), and d) display-dependent (habitat transitions depending on aerial display state). We fitted each model using maximum-likelihood estimates and extracted their AIC value. We predicted that the habitat-dependent model would be the best in the model set to explain our data, as habitat should influence the probability of gaining or losing aerial display, but exhibiting or not aerial displays should not influence transitions between habitats. Indeed, the habitat-dependent model had the lowest AIC in 98.3% of the trees (max. ΔAIC = 1.50; Figure S2). However, the weight of evidence in its favour was not unequivocal^41^ (*i.e.*, Akaike weight, *w*_*i*_ ≤ 0.90) in 92.5% of the trees (median = 0.84; range = 0.32 to 1.00), with the correlated model having a weight of evidence as high as 0.68 (median = 0.15; Figure S3). For this reason, and to avoid having to use subjective ΔAIC thresholds to select a single best model, we decided to perform model averaging of the estimates. We did that by weighting the transition rates estimated in each of the four models by the model’s Akaike weight, resulting in a single model-averaged estimate of each transition rate per tree (Figure S4). We then evaluated these transition rate estimates to evaluate whether gains of aerial display are more likely in open habitats than in forests. Across the 1,000 trees, we found that transition rates to aerial displays are 7.03 ± 1.28 (median ± SD) times more likely in open habitats than in forests, as predicted by our hypothesis (Figures 2, S5).

**Figure 2.**
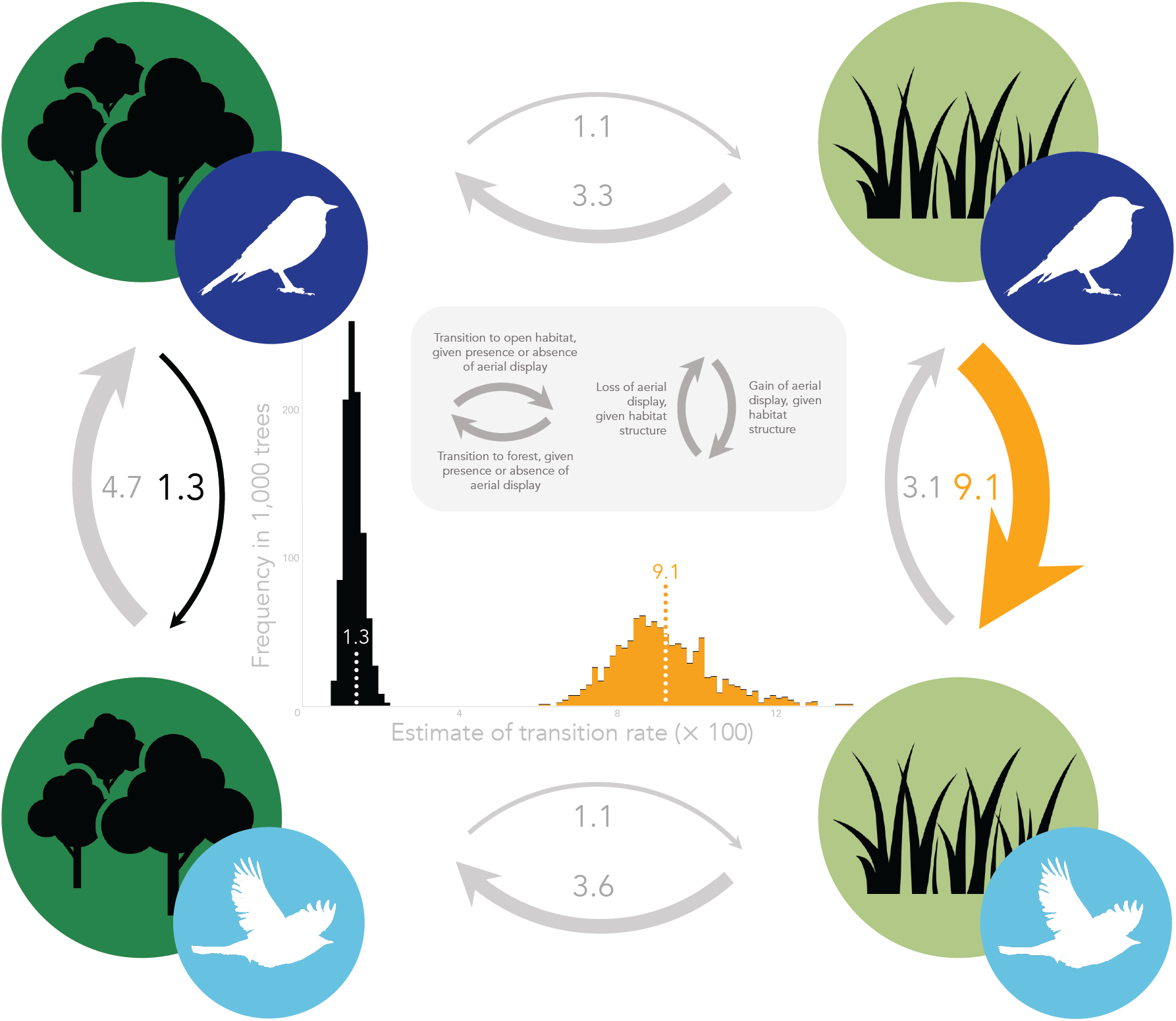
Evolutionary transition rates between habitat structure and aerial display. Arrows are weighted by the values beside them, which indicate the median (× 10^2^) model-averaged maximum-likelihood estimate of each rate across 1,000 different trees. Median transition rates to aerial displays are approximately seven times higher in open habitats (orange arrow) than in forests (black arrow; see Figure S5 for the distribution of orange rate to black rate ratios). In the centre, distribution of estimates of the two rates of interest across the 1,000 trees (see Figure S4 for the distribution of estimates of all transition rates).

## Discussion

Our findings support the hypothesis that aerial displays are more likely to evolve in open-habitat passerines than forest ones. This macroevolutionary pattern is likely the result of physical properties of the environment that allow aerial displays to transmit more broadly and effectively in open habitats than forests. Therefore, our order-level analysis of 469 species provides the most comprehensive evidence of sensory drive acting on signal evolution to date, as far as we are aware. It is also the first phylogenetic analysis to find evidence of habitat shaping the evolution of gestural displays. Previous studies had recognized the role of habitat structure in shaping sound-, colour-and ornament-based signals in lizards and birds^42–47^, but failed to detect an influence of habitat on the structure of motion-based signals (*e.g.,* in fiddler crabs^16^). More than seven decades after Armstrong’s^34^(p. 247) remark that “high-flying […] displays are most characteristic of birds of the open country”, we offer compelling evidence of the evolutionary process that likely underlies this pattern. Armstrong then goes on to state that the “prime consideration [of display-flights] is to attract the attention of the immigrant females”. Indeed, all displays analysed in our study likewise have a sexual function. Therefore, the fact that these are sexual displays allows us to discuss how our findings about sensory drive may interact with sexual selection, particularly regarding its direction and intensity.

Classical sexual selection models such as Fisherian runaway^48–50^and handicap^51,52^offer little prediction about which sexual traits should evolve and to what direction^53^. The handicap model, for instance, predicts how costly a signal should be, but not how these costs should be expressed^54^. The Fisherian runaway process assumes an arbitrary direction of evolution altogether^55,56^. Passerine gestural displays may seem particularly unpredictable, with multiple, clade-specific evolutionary forces acting on their evolution^28,57,58^. Yet, here we show that a simple ecological factor, habitat structure, predicts to a great degree the presence of a major gestural component. In the paper introducing the sensory drive hypothesis, Endler^5^highlighted an important aspect of sensory drive: that it allows us to predict which specific sexual traits should evolve as signals and to what direction. Our findings provide an emblematic example of how the signalling environment may help predict the direction of evolution of sexual signals.

Our system also allows us to explore the interplay between sensory drive and sexual selection from a different perspective. The effectiveness of aerial displays is influenced by the habitat because the signal can reach a broader range in open habitats than in forests. For males, this means that a male performing aerial display is able reach a higher number of females in open habitats than in forests. From the female perspective, it means that females are able to assess more males per unit of time in open, unobstructed habitats, thus increasing the number of potential mates they can sample during the mate choice process. Recent comparative and simulation data show that when females can sample more males during mate choice, sexual selection is stronger and, consequently, male ornaments are expected to be more extreme^59^. Thus, besides driving the evolution of aerial displays through sensory drive, we suggest that open habitats may intensify sexual selection on male aerial displayers by enabling females to sample more males in the population.

## Methods

### Study group

Passeriformes is, by far, the most speciose order of birds, encompassing 5,966 species^37^that show a diverse array of sexual behaviours and habitat-selection strategies^31^. Such diversity, and the fact that Passeriformes have well-sampled phylogenies^37^, make them an appropriate group for the purposes of this study. We searched for information on all extant species that regularly breed in the New World, excluding oceanic-island endemics (*e.g.*, Hawaii, Galápagos and Juan Fernández). According to the American Ornithologists’ Union^60–62^, 2,393 species meet these criteria, but an additional 49 species are recognized by Jetz et al.^37^, totalling 2,442 targeted species (Supplementary Data). For each species, we answered the following questions, which brought forth the three binary variables used in the analyses: (i) do individuals perform aerial display?; (ii) do individuals occupy primarily open or forest habitats?; and, in the case of forest species, (iii) do individuals forage exclusively in canopy stratum? Each of these variables will be detailed below.

### Display data

We collected display data from a variety of sources: life-history databases (*Handbook of the Birds of the World Alive*^31^, accessed 19 December 2016–10 March 2018; Cornell Lab of Ornithology’s *Neotropical Birds Online*^29^, accessed 21 August 2017–10 March 2018), natural-history books (A. C. Bent’s *Life histories of North American birds* series^63–72^; A. F. Skutch’s *Life histories of Central American birds*^73–75^; A. Wetmore’s *The birds of the republic of Panama*^76,77^; H. Sick’s *Ornitologia brasileira*^35^), an online media database (HBW’s *Internet Bird Collection*, http://hbw.com/ibc, accessed 4–10 October 2017) and a data paper^78^(see Supplementary Methods for details).

For data collection purposes, we considered sexual display any ritualized, sexually selected gesture or posture. The question *do individuals perform aerial display?* was answered *yes* (1) for a species only if any of the data sources claimed that individuals perform a sexual display that includes a flight component (*i.e.,* an aerial display). Otherwise, it was answered *no* (0) if any source claimed either (a) that individuals do not perform aerial sexual display, or (b) that they perform sexual display that does not include a flight component. Note that, for this question to be answered – be it with a yes or a no – a sexual display must have been mentioned by at least one data source (lack of any mention would result in NA). We used a set of criteria to infer from the available information whether a display is sexual (*i.e.*, sexually selected) and thus relevant for data collection purposes (Table 1). For some species (*N* = 29), information was ambiguous about whether the mentioned displays were sexual (see Table 1 for what we considered ambiguous). We excluded the ambiguous species from the analyses in the main text, but we also ran a phylogenetic logistic regression including them to assess whether it would change the results; it did not (see Supplementary Results).

**Table 1.**
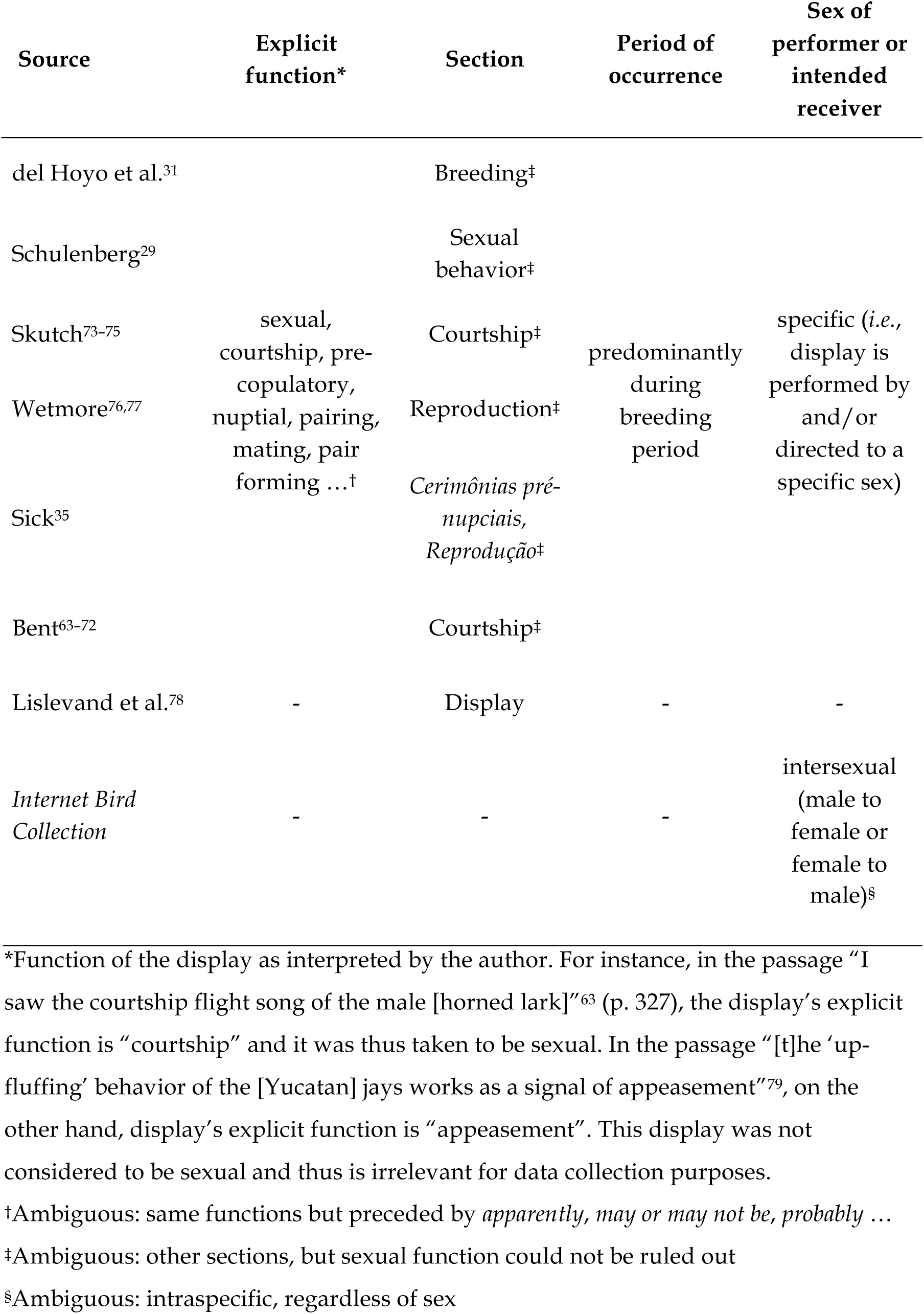
Criteria used to infer whether a display described in the literature is sexual (*i.e.*, sexually selected) or not. For a display described in a given data source, if the available textual information matched any corresponding column below, display was considered to be sexual. For instance, for a display described in del Hoyo et al.^31^, IF explicit function is sexual, courtship, … OR section is Breeding OR period of occurrence is predominantly during breeding period OR sex of performer or intended receiver is specific, THEN display was considered to be sexual. Footnotes refer to criteria that indicate that display may or may not be sexual; we repeated analyses including or excluding such displays (see Supplementary Results).

We defined flight component as the flapping of wings while the individual is not perched. In our conception, aerial displays are only likely to be more effective in open habitats if the flight is high enough that the bird surpasses the open vegetation. In spite of that, we decided to adopt an inclusive definition of flight component – one that includes even flights with little or no vertical gain – because for most species there was no information about the height of flight during display. This decision is likely conservative because most of the species with horizontal-flight displays we know of are forest dwellers (*e.g., Conopophaga, Rupicola*, many Pipridae, *Mionectes, Platyrinchus*). For some species (*N* = 9), we were unable to determine with certainty whether the sexual display included or not a flight component. This occurred when we came across ambiguous descriptions such as “occasional jumps/leaps into the air”^78^, “flitting from twig to twig”^75^(p. 45), and “move restlessly from branch to branch” (Mitchell 1957 *in* ^29^). We dealt with such uncertainty by repeating analyses considering flight component either absent (main results) or present (Supplementary Results) in the display; the results were qualitatively similar.

Overall, we were able to determine whether 470–499 passerine species (depending on how strict we were in treating a display as sexual) exhibited aerial displays or not, which represents 19.2–20.4% of the initial sample set. The remaining species were classified as NA and thus dropped from the analyses (this same decision was applied to the other variables as well).

### Habitat data

We collected data about habitat structure (for all passerines) and foraging stratum (for forest passerines only) from Parker et al.^80^and del Hoyo et al.^31^. The question *do individuals occupy primarily open or forest habitats?* was answered *open* (1) if first (primary) habitat listed for the species in Parker et al.^80^was of type *non-forest* or *aquatic*, and *forest* (0) if primary habitat was of type *forest* (see Supplementary Methods for which habitats are classified as non-forest, aquatic and forest by Parker et al.^80^). We were able to collect habitat data for 2,160 species (88.4% of the species in our initial sample set) from this source, which only lists birds breeding in the Neotropical region (*i.e.*, “from northern Mexico to the southern tip of Argentina, including the West Indies”^80^, p. 120). For the remaining species, we assigned the first habitat listed in section *Habitat* in del Hoyo et al.^31^to the closest habitat category used by Parker et al.^80^and determined whether it was a non-forest-, aquatic-or forest-type habitat. In a random sample of 100 species evaluated from both sources, agreement as to openness of habitat was very high (97%; posterior mean correlation = 0.97; 95% CI: 0.88 to 1). We were able to determine primary habitat for 2,436 species (99.8% of the initial sample set).

We searched for information about stratum only for species that occupy forest-type primary habitats (*N* = 1,864 species). The question *do individuals forage exclusively in canopy stratum?* was answered *yes* (1) if all foraging strata listed for the species in Parker et al. (1996) were either *canopy* or *aerial*, and *no* (0) if any other forest stratum was listed. For species not listed in Parker et al. (1996), we referred to del Hoyo et al. (2018) using the same criteria (but considering *subcanopy* as *canopy*). We were able to determine foraging stratum for 1,709 species (91.7% of forest species). Stratum data were only used in the analysis whose results are shown in the Supplementary Information.

### Phylogeny

We used species-level Passeriformes trees based on the Hackett backbone from Jetz et al.^37^(downloaded from http://birdtree.org on 18 March 2018). In each analysis, we pruned trees to match the corresponding species for which we had complete data (*i.e.*, species with missing data were dropped from the trees). Of the 469 species that were included in the main analyses, 415 (88.5%) have had their phylogenetic position determined based on genetic data^37^(Supplementary Data).

### Phylogenetic logistic regressions

We used phylogenetic logistic regressions^38^to test for the influence of primary habitat structure on the probability of passerines exhibiting aerial display (*N* = 469 species), and to test for the influence of foraging stratum on the probability of forest passerines exhibiting aerial display (*N* = 235 species; Supplementary Results). In the first analysis, we compared a model with no predictor variable to one including habitat structure as a categorical predictor. We fitted each model using the function phyloglm from package phylolm^81^(version 2.5) in R^82^(version 3.4.3). To account for phylogenetic uncertainty^83,84^, we fitted each model iteratively using a sample of 1,000 different trees from Jetz et al.^37^, totalling 2,000 models. We fitted null models manually, and models predicted by habitat using the package sensiPhy^85^(version 0.8.1). From each of the 2,000 models, we extracted Akaike information criterion (AIC) value as well as the coefficients’ estimates and *p*-values. The model with the lowest AIC was selected in each iteration, as long as the competing model’s ΔAIC was greater than 2. Estimates were considered statistically clear if *p* ≤ 0.05^86^. According to our hypothesis, we expected that the model containing habitat structure as a predictor would be selected in most iterations, with a positive slope estimate.

### Correlated evolution of discrete characters

A different approach to test the hypothesis that aerial displays are more likely to evolve in open-habitat passerines than forest ones was to use Pagel’s^39^method of detecting correlated evolution between characters. We performed this analysis using the Discrete module in BayesTraits V3.0.1^40^under a maximum-likelihood (ML) approach. In order to test our hypothesis with this method, we had to ultimately assess whether the evolutionary rate of gain of aerial displays is higher in species that live in open habitats than in species that occupy forests. But a few steps precede this assessment, and we will get to them below.

Pagel’s method is based on continuous-time Markov chains with finite state space (Mk model), which calculates the probability that a trait will change from state *i* to state *j* over a very (infinitesimally) short time interval^39,87^. This probability is called instantaneous transition rate and is represented by the parameter *q*_*ij*_. In this study, we have two binary characters that, combined, result in four possible states a species may assume: forest without aerial display, open habitat without aerial display, forest with aerial display, and open habitat with aerial display. Let us call these states 1 to 4, respectively – because it is the order in which they appear in Figure 2. In this figure, each arrow corresponds to a parameter *q*. By looking at *q*_12_, for example, we are looking at the probability that a species that does not exhibit aerial display transitions from forest to open habitat. Note that there are eight possible parameters (eight arrows) because the diagonals are assumed to be zero; since we are dealing with a very short time interval, it is assumed that both traits cannot change at the same time.

Pagel^39^suggests four biologically plausible models by which two characters can evolve. If the characters evolve independently (*i.e.*, independent model), the transition rates in one character do not depend on the state of the other character. In this model, the transition from forest to open habitat, for example, is assumed to be the same regardless of whether the species does or does not exhibit aerial display (*q*_24_ = *q*_12_).Likewise, the transition from presence to absence of aerial display is the same regardless of the species’ preference of habitat structure (*q*_31_ = *q*_43_). The same holds true for the remaining two pairs of parameters (*q*_13_ = *q*_34_; *q*_42_ = *q*_21_). The independent model is the most parsimonious one, because it has only four free parameters (with the other four restricted to have the same value as the free ones). Alternatively, if the characters evolve in a correlated manner (*i.e.*, correlated model), all eight parameters are free to vary: gain of aerial display depends on habitat (*q*_12_ ≠ *q*_24_), transition from forest to open habitat depends on presence of aerial display (*q*_13_ ≠ *q*_34_), and so on. This is the least parsimonious model, with eight free parameters. In between the independent and correlated models, Pagel suggests that one character may influence the evolution of another, but not the other way around. Thus, if only presence/absence of aerial display influences preference for habitat structure (*i.e.*, display-dependent model), the transition rates between habitats should be different depending on whether the species exhibits or not aerial display (*q*_12_ ≠ *q*_34_; *q*_21_ ≠ *q*_43_), but gain and loss of aerial display are assumed to be the same regardless of the habitat (*q*_13_ = *q*_24_; *q*_31_ = *q*_42_). Analogously, in the habitat-dependent model, gain and loss of aerial display are different according to the habitat (*q*_13_ ≠ *q*_24_; *q*_31_ ≠ *q*_42_), but transitions between habitats are the same regardless of aerial display presence (*q*_12_ ≠ *q*_34_; *q*_21_ ≠ *q*_43_). Both habitat-dependent and display-dependent model have six parameters each.

We extracted the AIC value of each model using the following equation, where *k* is the number of parameters (*i.e.*, number of free *qs*) and 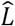 is the maximum likelihood:

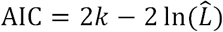

We also calculated the Akaike weights (*w*_*i*_.) of each model to assess the weight of evidence in its favour relative to all the other models in the set^41,88^. Next, we used a model-averaging approach to estimate the parameter values. For a given parameter *q*, we did that by weighting the ML estimate in each of the four models 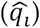 by the model’s *w*_*i*_., resulting in the model-averaged ML estimate 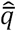^41,89^:

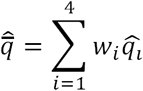

We decided to use a model-averaging procedure because it offers advantages such as basing inference not just on the one model estimated to be the best, but on all *a priori* models^89^, and avoiding the need for subjective thresholds to choose which model is the best^90^. After performing model averaging, we were then able to properly test our hypothesis by comparing the model-averaged ML estimate of the rate of gain of aerial display in open habitats (*q*_24_) and forests (*q*_13_), predicting that *q*_24_ > *q*_13_.

All steps described above were repeated for each tree in the 1,000-tree sample from Jetz et al.^37^to account for phylogenetic uncertainty, and the rates shown by the arrows in Figure 2 are the median model-averaged ML estimate for each transition rate, multiplied by 10^2^, across the 1,000 trees.

## Supporting information

## Data availability

All of the R code and data used in our analyses will be made available on Dryad upon acceptance of the paper.

## Acknowledgments

We thank R. Maia and L. Manica for critical input throughout the development of the project; D. Caetano and D. Muniz for comments and suggestions on earlier drafts of the manuscript; D. Caetano, A. Meade, G. Burin, and C. Botero for help with analyses; P. Guimarães Jr. for help with data collection; J. Quental, K. Chamberlain and K. Hay for permission to use photos (and G. Smith for licensing under CC). Lastly, we thank the many naturalists and ornithologists out in the field watching birds as they dance; their observations are the basis of this study. This study was financed in part by the Coordenação de Aperfeiçoamento de Pessoal de Nível Superior – Brasil (CAPES) – Finance Code 001.

## Competing interests

The authors declare no competing interests.

