## Supplementary material for "Habitat structure drives the evolution of aerial displays in birds"

**Supplementary Methods**

*Display data collection*

We used different types of references as sources for display data, including books, online databases and a data paper. For each of these sources, we adopted different strategies to optimize the search for relevant display information (Table S1).

**Table S1.** References used as sources for display information, number of relevant species comprehended by them, strategy used to search for relevant information, and dates of access (for online resources only).

| Reference | # relevant species* | Search strategy | Dates accessed |
|---|---|---|---|
| del Hoyo et al.[1] | 2,418, in individual species accounts | in each species account, read sections *Voice* and *Breeding*, and look up keyword *display* | Dec 2016, Jan 2017, Mar–Jun 2017, Aug 2017, Jan–Mar 2018 |
| Schulenberg[2] | 561, in individual species accounts | in each species account, read section *Behaviour* | Aug 2017, Mar 2018 |
| Bent[3–12] | 306, in ten books | in each book, look up keywords *court*, display, aerial, flight song, song flight* | - |
| Skutch[13–15] | 149, in three books | in each book, look up keywords *court*, display, aerial, flight song, song flight* | - |
| Wetmore[16,17] | 462, in two books | in each book, look up keywords *court*, display, aerial, flight song, song flight* | - |
| Sick[18] | 920 | look up keywords | - |

| | | | |
|---|---|---|---|
| | | *display, demonstraç*, exib*, cerim*, pulo-voo, corte, ritual* | |
| *Internet Bird Collection* | n/a (1,878 videos analysed) | search keywords *display, song flight, aerial* OR *courtship* | Oct 2017 |
| Lislevand et al.[19] | 405 | extract information from column *Display* | - |

*Number of species comprehended in data source that are relevant for the purposes of this study. For Schulenberg[2], it is the number of accounts that have credited authors — which we took as indication that available information for the species had been exhaustively searched. For del Hoyo et al.[1], it is the number of full accounts — *i.e.*, whose sections were all filled in.

*Habitat data classification*

Our main source of information about habitat preference was Parker et al.[20]. Primary habitats categorized by them as non-forest or aquatic were included in our dataset as *open*, whereas forest-type habitats were accordingly considered as *forest* in our dataset. Table S2 contains all habitats listed by Parker et al.[20].

**Table S2.** Classification of habitats in Parker et al.[20] and our study.

| Original classification | Our classification | Habitat |
|---|---|---|
| Forest* | Forest* | tropical lowland evergreen forest; flooded tropical evergreen forest; river-edge forest; montane evergreen forest; elfin forest; *Polylepis* woodland; tropical deciduous forest; gallery forest; southern temperate forest; pine forest; pine-oak forest; white sand forest; palm forest; mangrove forest; secondary forest |
| Non-forest | Open | arid lowland scrub; arid montane scrub; semihumid/humid montane scrub; *cerrado; campo* grasslands; low, seasonally wet grassland; |

| | | |
|---|---|---|
| | | southern temperate grassland; northern temperate grassland; *puna* grassland; *páramo* grassland; riparian thickets; river island scrub; pastures/agricultural lands; second-growth scrub |
| Aquatic | Open | freshwater marshes; saltwater/brackish marshes; coastal sand beaches/mudflats; coastal rocky beaches; riverine sand beaches; freshwater lakes and ponds; alkaline lakes; rivers; streams; bogs; coastal waters; pelagic waters |

*Includes forest edges.

*PLogReg: is our missing data phylogenetically biased?*

An elevated percentage of missing data such as we had in this study (80.8%) may bias the results if the absence of data is not independent of phylogeny, which would compromise our ability to extrapolate the results to all New World passerines. To assess whether the absence of data is phylogenetically clumped, we tested for phylogenetic signal in the missing display data using the function miss.phylo.d from the package sensiPhy[21] (version 0.8.1), which calculates the $D$ statistic[22,23] for missing data. A $D$ value of 0 means missing data is as phylogenetically conserved as if it had evolved under Brownian movement, while a value of 1 would emerge if missing data were randomly shuffled relative to the tips of the phylogeny. Mean $D$ statistic for missing display data in 50 randomly sampled trees corresponding to the initial sample set (2,442 species) was of 0.45 (± 0.01 SD), clearly ($p < 0.001$) different from both Brownian motion ($D = 0$) and random ($D = 1$) models in all trees.

*Maddison and FitzJohn's[24] criticism of Pagel's method*

Pagel's correlation method[25] has been found to report a significant correlation between traits even from evidence that may well be coincidental[24]. Such evidence comes from evolutionary scenarios in which one or both traits have a low number of transitions and/or when transitions are concentrated within a single lineage (Figures 1c,d in Maddison and FitzJohn[24]). To assess how robust our data is to the concerns raised by Maddison and FitzJohn[24], we estimated the number of character changes for both our traits using stochastic mapping (SIMMAP[26,27]) conditioned on the ML-estimated, model-averaged transition rates (main results, Figure S4) for each of 100 trees. Median number of changes was 109 (range: 92 to 126) for habitat and 191 (range: 149 to 244) for display. The ancestral state estimation of our characters shows that these transitions are dispersed throughout the phylogeny, rather than concentrated in a single clade (Figure S1). The fact that our traits have multiple, dispersed transitions diminishes the chances that our results suffer from the issues raised by Maddison and FitzJohn[24]. Nevertheless, studies have not shown how many transitions are enough, and how dispersed they must be, in order for results to be considered free from these issues; hence the importance of using complimentary analyses, as we did with Pagel's and PLogReg[28] (main results).

**Supplementary Results**

*PLogReg: does relaxing the definition of sexual display change the results?*

As we explained in the Methods section, for a species to be considered in our analyses, a sexual display had to be mentioned in at least one data source. For 29 species, we were not able to determine whether the mentioned display was or was not sexual. We excluded these species from the PLogReg whose results are shown in the main text. However, we also ran this analysis including them to assess whether there was a significant change in results. In practice, this means treating as 0 (*does not exhibit aerial display*) or 1 (*exhibits aerial display*), 29 species that were previously considered NA (missing data), resulting in a sample of 498 species. The model containing habitat structure as predictor was selected in all of the 1,000 iterations, with a mean ΔAIC of 18.89 (min = 8.09, max = 41.91). Mean slope estimate for open habitat was 0.517 (95% CI: 0.514 to 0.520), with a mean *p*-value of 0.015 (≤ 0.05 in 99.6% of iterations). Relaxing the definition of sexual display, thus, provides stronger evidence in favour of the hypothesis that aerial displays are more likely to evolve in open-habitat passerines.

*PLogReg: does relaxing the definition of flight component change the results?*

We also mentioned in the Methods that we were not able to determine whether the sexual display by nine species included or not a flight component. The PLogReg results in the main text consider these species as not exhibiting aerial display (0), but we repeated the analysis considering them as exhibiting aerial display (1). The sample size is the same as in the main analysis (*N* = 469 species). The model containing habitat structure as a predictor was selected in all of the 1,000 iterations, with a mean ΔAIC of 26.46 (min = 11.52, max = 53.12). Mean slope estimate for open habitat was 0.592 (95% CI: 0.589 to 0.595), with a mean *p*-value of 0.007 (≤ 0.05 in 100% of iterations). Relaxing the definition of flight component, thus, provides even stronger evidence in favour of the hypothesis that aerial displays are more likely to evolve in open-habitat passerines.

*Does a Bayesian phylogenetic logistic regression yield different results?*

In simulations[29], PLogReg showed slightly inflated Type I errors when the independent variable evolves under Brownian motion and the species sample is relatively large ($32 \leq N \leq 256$). The *D* value[22] estimated for our independent variable — habitat structure — using 50 randomly sampled trees is of 0.13 ± 0.03 (mean ± SD), and not clearly different from 0 (Brownian motion) in any tree[23]. This means that the null hypothesis — that aerial displays are equally likely to evolve regardless of habitat structure — may have been rejected slightly more often than it should in our PLogReg result. To ensure that the positive, mostly clear, value of open habitat predicting aerial display is not a by-product of Type I error, we also performed a phylogenetic logistic regression using a Bayesian approach (Stan[30]) with the package brms[31] (version 2.1.0). We ran two MCMC chains for 2,000 iterations (after a burn-in of 2,000) for each of 50 randomly sampled trees, and extracted the mean and 95% confidence interval from the total posterior distribution of 200,000 estimates of slope (*i.e.*, 2,000 iterations × 2 chains × 50 trees = 200,000). The posterior mean estimate for slope was positive and higher than estimated by the PLogReg ($ß_{habitat}$ = 2.020), although the 95% confidence interval does overlap zero (-1.532 to 7.818). In the simulations by Ives and Garland[29], another Bayesian method to test phylogenetic logistic regressions — MCMCglmm[32] — had the opposite effect of PLogReg: it rejected the null hypothesis (*i.e.*, excluded 0 from the 95% CI) *less* often than it should. The fact that the confidence interval of the MCMC posterior distribution overlaps zero may thus be partly due to Type II error.

*Testing the influence of forest stratum on aerial display*

Because of its physical properties, we expected the canopy stratum of forests to have a similar effect on aerial display effectiveness as open habitats do. Like in open habitats, flying above the canopy during the display should put the signaller under more intense light and reduce the vegetation obstructing the signaller from the perspective of potential receivers. Canopy aerial displayers should also put themselves against the same uniform background open-habitat aerial displayers do, the sky. Thus, we ran a PLogReg to evaluate whether canopy stratum influences the probability that a forest species exhibits aerial display. Our sample size for this test was of 234 species (*i.e.*, forest species for which we hadn data about stratum and display), and we used the same sample of 1,000 trees from Jetz et al.[33]. The model that included stratum as a predictor variable was selected ($\Delta$AIC > 2) in 44.2% of the iterations. Despite having a small negative value ($\beta_{canopy}$ = -0.091; 95% CI: -0.103 to -0.079), the slope estimate for stratum was statistically unclear in all iterations (mean *p*-value = 0.549), thus offering little support to our prediction that canopy passerines are more likely to perform aerial display than passerines that use lower strata. This result contrasts with previous phylogenetic analyses that showed an influence of forest stratum on avian gestural displays, plumage coloration and vocalization[34–36].

### Supplementary Figures

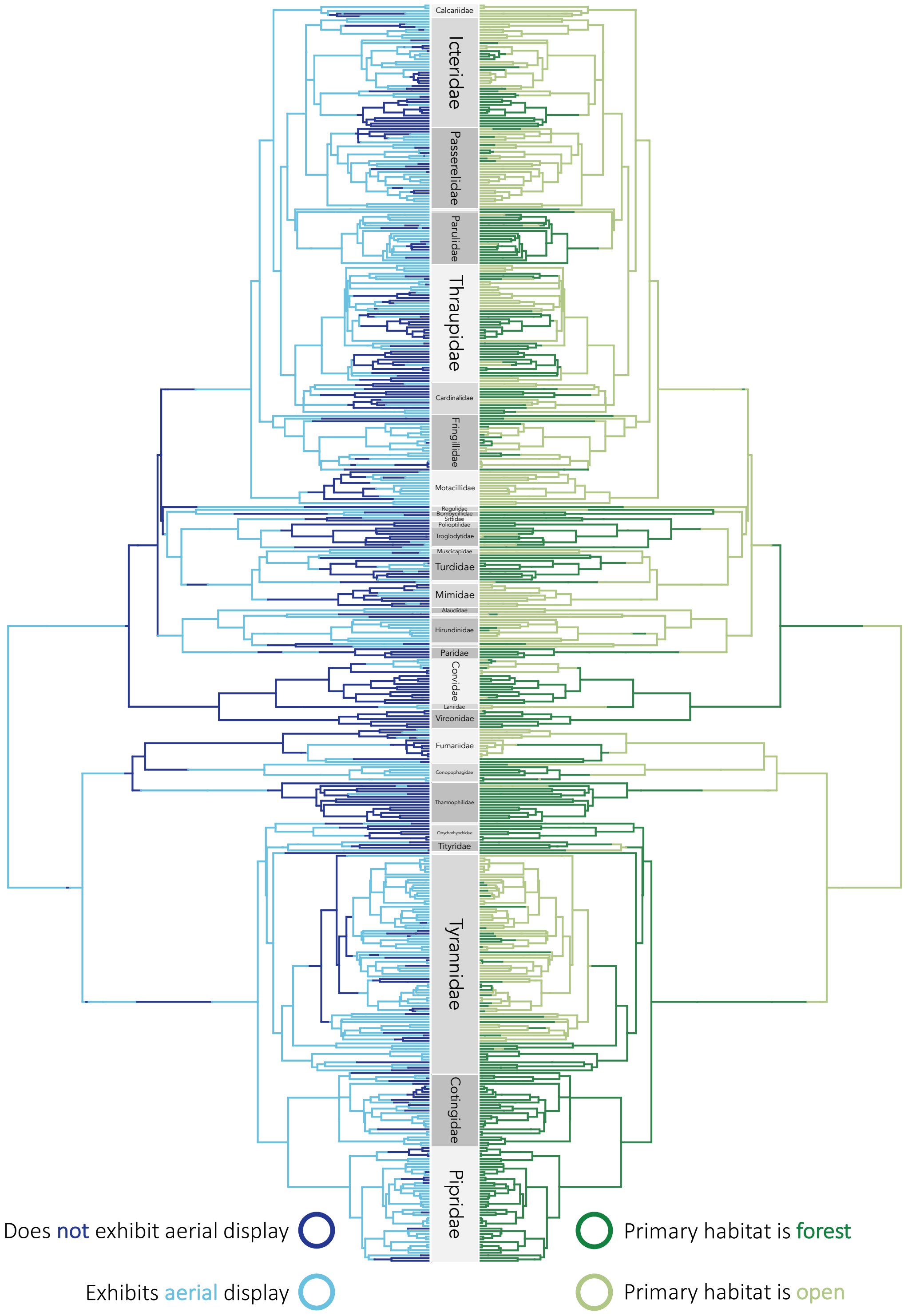


**Figure S1**. Ancestral state estimation of aerial display and habitat structure, using stochastic mapping (SIMMAP) conditioned on the model-averaged, maximumlikelihood estimates of transition rates (see Results) and using the same tree as in Figure 1 in the main text.

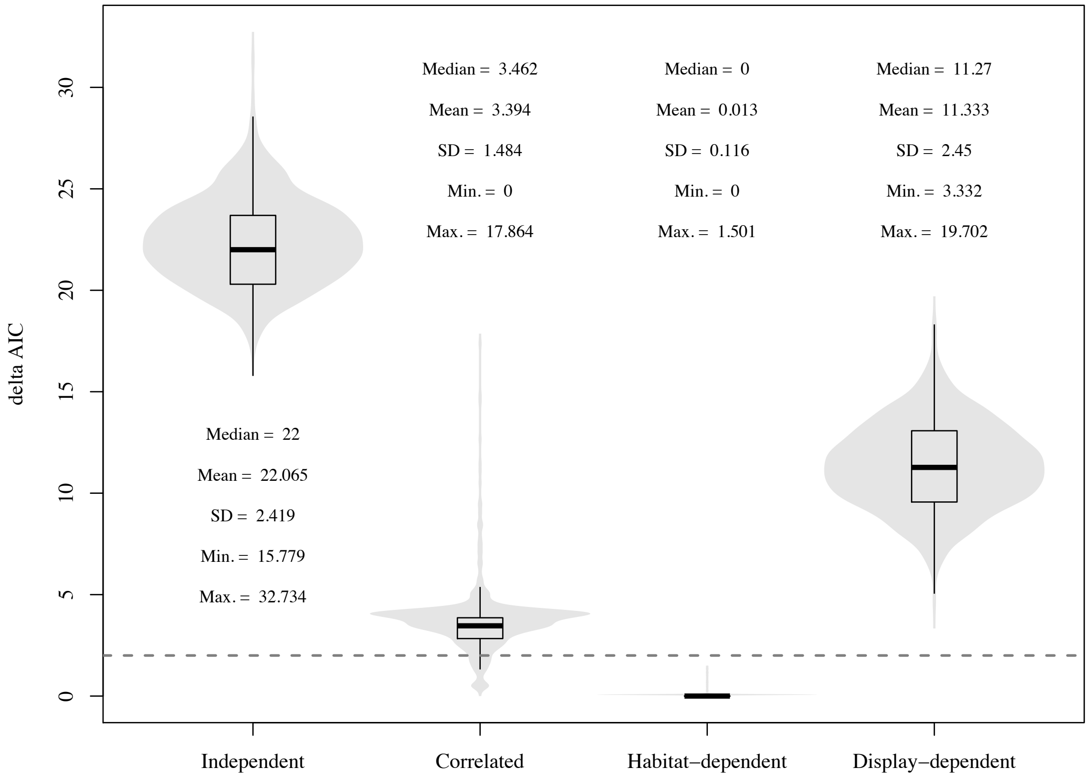


**Figure S2.** Violin plots and boxplots of the ΔAIC values of each model across 1,000 trees. Dashed line represents commonly used threshold under which models are considered competitive (ΔAIC < 2). Note that we did not use that threshold in our study, instead adopting a model-averaging approach (see Methods in the main text). In boxplots, band represents the median, box limits represent the interquartile range (IQR) and whiskers extend to the most extreme data within 1.5 IQR of the box; outliers are not plotted.

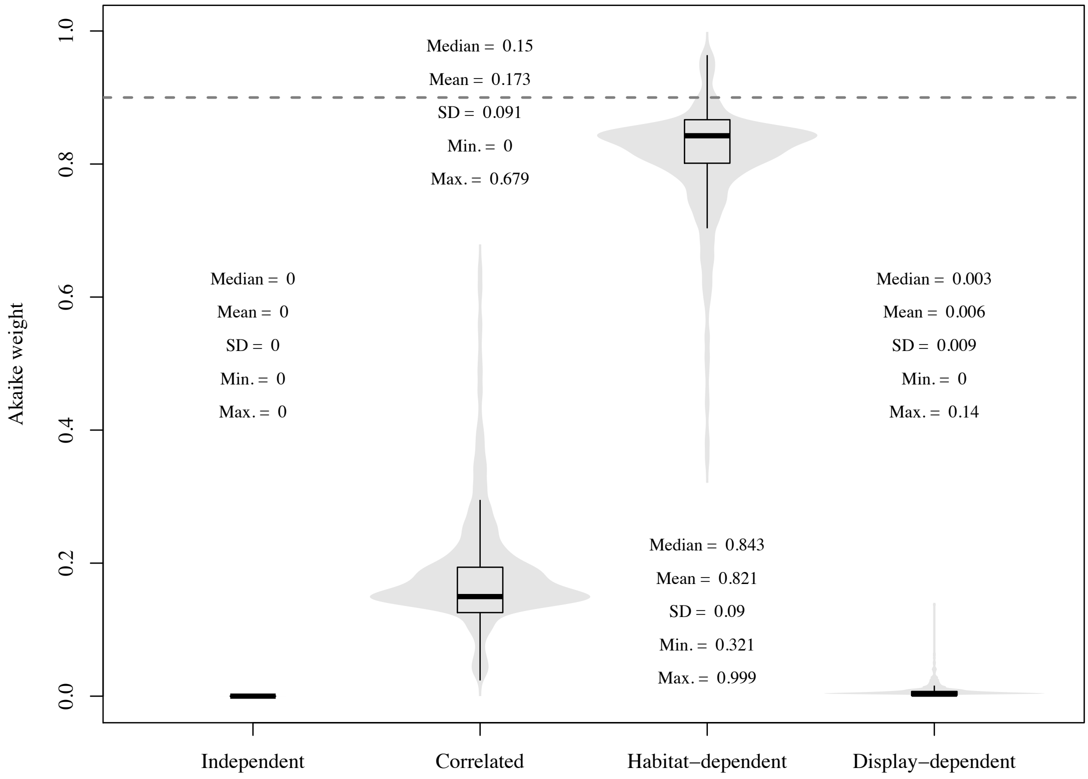


**Figure S3.** Violin plots and boxplots of the Akaike weights of each model across 1,000 trees. Dashed line represents threshold above which weight of evidence is usually considered unequivocal ($w_i \geq 0.9$). In boxplots, band represents the median, box limits represent the interquartile range (IQR) and whiskers extend to the most extreme data within 1.5 IQR of the box; outliers are not plotted.

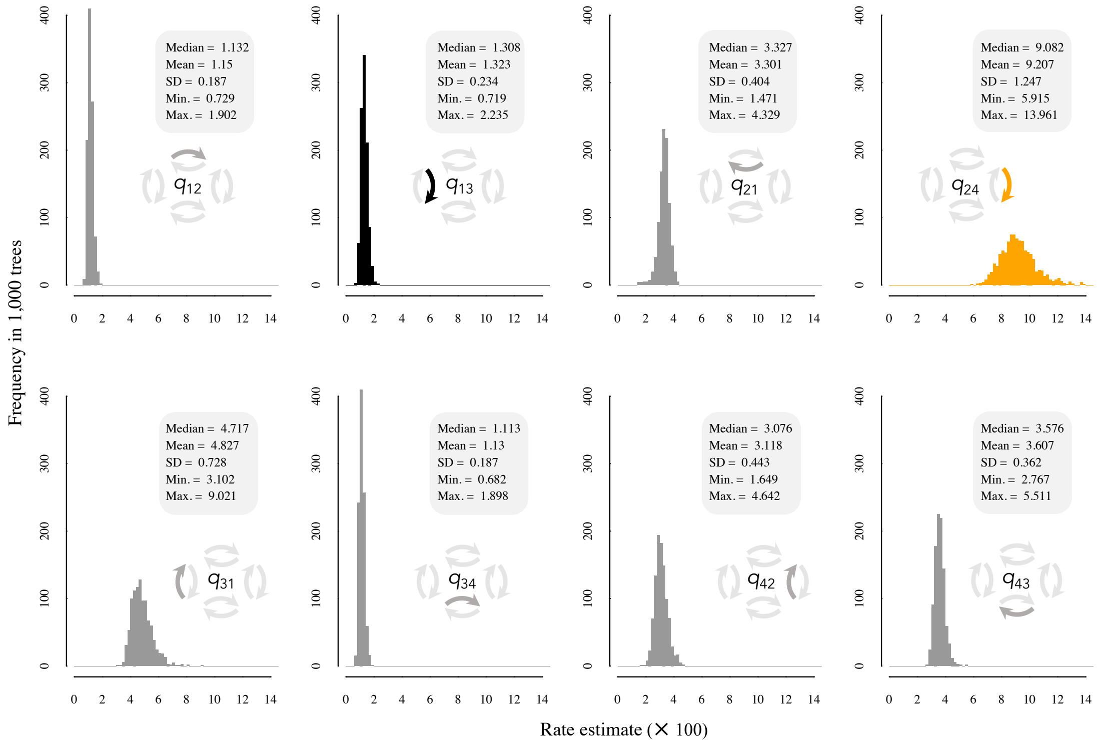


**Figure S4.** Distribution of the estimates of all transition rates across 1,000 trees. Arrow positioning of transition rates is the same as in Figure 2 in the main text.

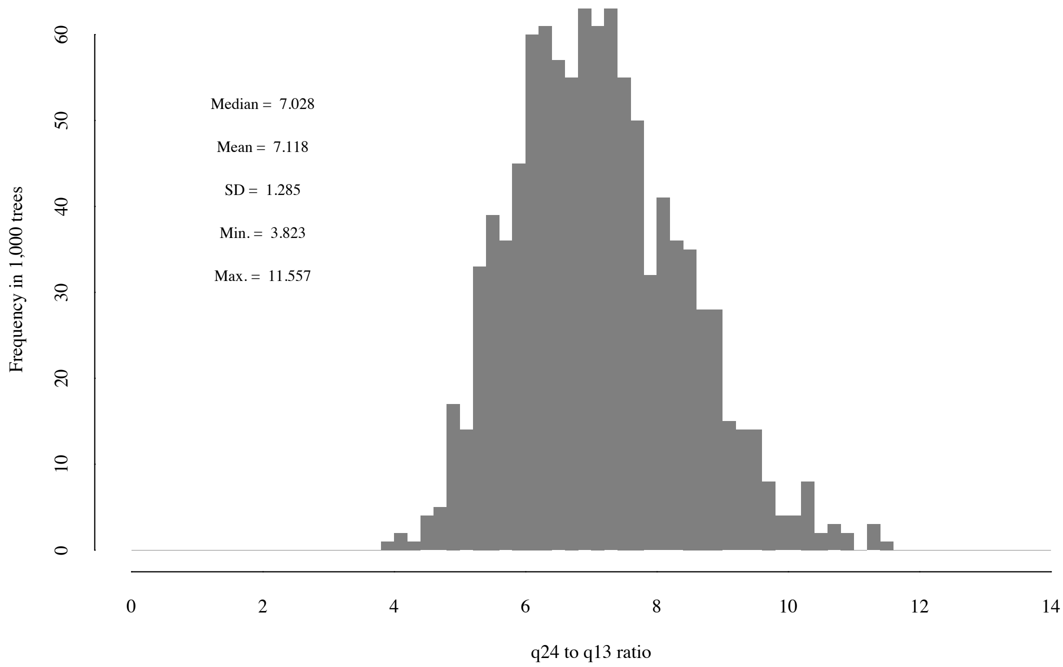

**Figure S5.** Distribution of $q_{24}$ (gain of aerial display in open habitats) to $q_{13}$ (gain of aerial display in forests) ratios across 1,000 trees. This ratio represents how many times more likely aerial display is to evolve in open-habitat species than in forest species.

20. Parker, T. A., III, Stotz, D. F. & Fitzpatrick, J. W. in *Neotropical birds: ecology and conservation* (eds. Stotz, D. F., Fitzpatrick, J. W., Parker, T. A., III & Moskovitz, D.

K.) 115–436 (Univ. of Chicago Press, Chicago, 1996).
